# Agent-based modeling of the prostate tumor microenvironment uncovers spatial tumor growth constraints and immunomodulatory properties

**DOI:** 10.1101/2023.07.27.550792

**Authors:** Maisa van Genderen, Jeroen Kneppers, Anniek Zaalberg, Elise Bekers, Andries M Bergman, Wilbert Zwart, Federica Eduati

## Abstract

Inhibiting androgen receptor (AR) signaling through androgen deprivation therapy (ADT) reduces prostate cancer (PCa) growth in virtually all patients, but response is temporary, and resistance inevitably develops, ultimately leading to lethal castration-resistant prostate cancer (CRPC). The tumor microenvironment (TME) plays an important role in the development and progression of PCa. In addition to tumor cells, TME-resident macrophages and fibroblasts express AR and are therefore also affected by ADT. However, the interplay of different TME cell types in the development of CRPC remains largely unexplored.

To understand the complex stochastic nature of cell-cell interactions, we created a PCa-specific agent-based model (PCABM) based on *in vitro* cell proliferation data. PCa cells, fibroblasts, “pro-inflammatory” M1-like and “pro-tumor” M2-like polarized macrophages are modeled as agents from a simple set of validated base assumptions. PCABM allows us to simulate the effect of ADT on the interplay between various prostate TME cell types. The resulting *in vitro* growth patterns mimic human PCa.

Our PCABM can effectively model hormonal perturbations by ADT, in which PCABM suggests that CRPC arises in clusters of resistant cells, as is observed in multifocal PCa. In addition, fibroblasts compete for cellular space in the TME while simultaneously creating niches for tumor cells to proliferate in. Finally, PCABM predicts that ADT has immunomodulatory effects on macrophages that may enhance tumor survival. Taken together, these results suggest that AR plays a critical role in the cellular interplay and stochastic interactions in the TME that influence tumor cell behavior and CRPC development.

## Introduction

Prostate cancer (PCa) is the second most common cancer in men worldwide, with 1.4 million new cases and over 370,000 deaths annually^1^. Androgen receptor (AR) signaling plays a pivotal role in PCa initiation and progression, motivating the development of several therapies targeting this hormone-driven transcription factor over the years^2–4^. However, despite an initial treatment response in most patients, resistance to ADT inevitably develops, resulting in lethal metastatic castration-resistant prostate cancer (CRPC). Therefore, the development of new therapies that effectively treat or even prevent CRPC is critical^5^.

Recently, multiple studies have shown that the tumor microenvironment (TME) plays a key role in the development and progression of PCa^6–10^. The prostate TME consists of a variety of non-malignant cells, including fibroblasts and macrophages^11–14^. Cells in the TME influence PCa cell growth through chemical and physical interactions between tumor- and stromal cells, through angiogenesis, immune suppression, extracellular matrix (ECM) remodeling and tumor invasion^9, 15–17^. Although fibroblasts are mostly quiescent in healthy tissues, in the TME fibroblasts are in a state reminiscent of wound healing and are referred to as cancer-associated fibroblasts (CAFs)^11, 18, 19^. Another dominant component of the prostate TME is macrophages, which are highly plastic cells that can polarize into a spectrum of phenotypes. Conventionally, two extreme polarizations of tumor-associated macrophages are recognized: classically activated pro-inflammatory (M1) macrophages and alternatively activated anti-inflammatory (M2) macrophages^20, 21^. In general, M1-macrophages are anti-tumorigenic leading to tumor cell death, whereas M2-like macrophages are pro-tumorigenic, promoting tumor growth. These phenotypically distinct macrophages have been hypothesized to have contrasting effects on tumor progression^22^. Importantly, specific macrophage subtypes have a prognostic value for PCa patients, suggesting that the relative contributions of these subtypes are related to patient outcome^23^.

Interestingly, AR expression is not restricted to PCa cells, but is also expressed and functional in cells of the prostate TME, including fibroblasts and macrophages^24^. Consequently, interactions between cells of the prostate TME could potentially be affected by androgens and thus by AR-targeted therapies, including ADT. However, studies on ADT altering TME cell interactions in the context of primary PCa and CRPC development are limited and present conflicting results. Low levels of AR in stromal tissues are associated with an earlier onset of PCa recurrence^7, 25^. Indeed, AR signaling in the stroma has been reported to play a protective role in PCa development, as low AR expression in the TME is associated with a high-grade tumor and poor clinical outcome^7^. Previously, we have shown that AR inhibition in CAFs triggers PCa cell migration via paracrine regulation of CCL2 and CXCL8, which may contribute to PCa invasiveness and metastasis^25^. Alternatively, infiltration of tumor-associated macrophages (TAMs) influences disease progression toward CRPC development after ADT^26–28^. AR signaling in macrophages activates TREM-1 signaling, which subsequently leads to the secretion of pro-inflammatory cytokines that support PCa cell line migration and invasion^29^. In addition, AR has been described as an enhancer of macrophage and monocyte differentiation^30, 31^. However, it is not fully understood how the combined interactions between TME cells contribute to CRPC development and what the role of ADT is in these interactions.

Recently, computational agent-based models (ABMs) have been used to describe the complex interplay between cancer cells and TME cells^32–34^ by modeling individual agents that perform stochastic actions, thereby creating complexity from a simple set of base cell actions. Previously, ABMs have been successfully applied to study tumor stem cell growth^35, 36^, tumor cell migration^37^, avascular tumor growth^38^, radiotherapy optimization^39^ and response to immunotherapy in colorectal cancer^40, 41^. Recently we developed an ABM to study prostate cancer onset, however this does not account for the effect of therapy on the prostate tumor microenvironment^42^.

In this study we generated a PCa-specific ABM (PCABM) which includes the interactions between tumor cells, fibroblasts, and macrophages in relation to hormonal therapy. The PCABM is informed by *in vitro* prostate TME co-culture growth data, using particle swarm optimization (PSO). PCABM simulations show that CRPC is multifocal and arises from clusters of resistant cells within the prostate TME. In addition, fibroblasts play an indispensable role in regulating spatial proliferative constraints while simultaneously providing a protective niche for tumor cells from the tumoricidal effect of pro-inflammatory macrophages.

Recently, we reported a genome-wide CRISPR screen in PCa cells co-cultured with pro-inflammatory macrophages where we identified AR as a critical regulator of macrophage-mediated killing^43^. These studies revealed AR as a genuine tumor-intrinsic immunomodulator, with hormone deprivation preventing tumor cell killing by M1 macrophages. Consistent with this study in cell line models, our PCABM confirms *in silico* that ADT exposes immunomodulatory effects in the prostate TME, impeding macrophage-mediated tumor cell killing in androgen-deprived conditions. Cumulatively, our *in silico* model faithfully phenocopies both the response of tumor cells to hormonal stimuli, as well as the impact of therapy thereon in relation to its microenvironment.

## Materials & Methods

### In vitro Cultures

#### Cell culture and M1- and M2 macrophage differentiation

The prostate cancer cell lines LNCaP (ATCC CRL-1740) and LNCaP-abl (ATCC CVCL-4793) the monocytic cell line THP-1 (ATCC TIB-202) and immortalized foreskin fibroblast BJ cell line (CRL-2522) were cultured in RPMI-1640 (Gibco) supplemented with 10% fetal bovine serum (FBS, Sigma) and 1% penicillin-streptomycin (P/S, Gibco). For hormonal related experiments all cells were cultured in RPMI 1640 supplemented with 5% Dextran Coated Charcoal (DCC, Sigma) stripped-serum and 1% P/S 3 days before to the start of the experiment. AR was induced with 10nM R1881 (Sigma) supplemented RPMI-DCC. Cell lines were kept at low passage and regularly tested mycoplasma negative. THP-1 cells were stimulated with either 100ng/mL (for M1 macrophages) or 50ng/mL (for M2 macrophages) of phorbol 12-myristate 13-acetate (PMA, Sigma) for 48h, followed by 24h in fresh 10%FBS-RPMI. M1-macrophages were differentiated by 24h stimulation of 10ng/mL lipopolysaccharide (LPS, Sigma) and 10ng/mL interferon-*γ* (IFN-*γ*, Peprotech), while M2-macrophages were differentiated by 72h stimulation with 20ng/mL IL-4 (Peprotech) and 20ng/mL IL-13 (Peprotech).

#### Lentiviral vector and transduction

Lentivirus was generated in HEK293T cells cultured in 10% FBS, 1% P/S supplemented DMEM (Gibco). To produce LNCaP-eGFP cells, HEK293T were transfected using polyethylenimine (PEI) with packaging constructs (pMDLg/pRRE, pRSV-Rev, pCMV-VSV-G, AddGene). Virus was harvested after 24h, filtered with a 0.22*µ*m filter (Millipore) and snap frozen in liquid nitrogen. LNCaP cells were infected at a MOI *>* 2 and selected with 2*µ*g/mL puromycin (Sigma) and checked for eGFP expression regularly.

#### Three cell type co-culture assays

For co-culture assays, LNCaP cells and BJ fibroblasts were cultured together with either M1- or M2-like macrophages (**Supplementary Figure S1**). Additionally, LNCaP cells were cultured with BJ fibroblasts, M1- or M2-like macrophages separately. Firstly, 3750 THP-1 cells were seeded in a 96-well plate (CELLSTAR plate, 96w, F, *ν*Clear, TC, PS, black, lid, Greiner) in 100*µ*L medium per well. THP-1 cells were differentiated towards M1- or M2-like macrophages following the above-mentioned protocol. LNCaP-eGFP cells were added to differentiated macrophages with or without BJ fibroblasts (4:1 ratio). To investigate the effect of different hormone conditions on LNCaP cell survival, all cells were cultured in 5% DCC and 1% PS RPMI-1640 and stimulated with either DMSO (vehicle) or 10nM R1881. Additionally, cells were individually stimulated with either DMSO or 10nM R1881 for 24h, washed and co-cultured subsequently. LNCaP-eGFP cell fluorescence and proliferation was measured using IncuCyte Zoom (Essen BioScience) for 7 days. BJ fibroblast proliferation was measured separately by IncuCyte Zoom phase-contrast analysis.

#### Hormone conditions, apoptosis and resistant cell assays

To validate PCABM predictions on ADT effects, 3750 THP-1 cells were differentiated into M1- and M2 macrophages as described earlier in 5% DCC, 1% PS RPMI-1640. M1- and M2 macrophages were subsequently stimulated with either DMSO (vehicle) or 10nM R1881 for 24 hours. LNCaP-eGFP cells were seeded at a density of 15000 cells per well in a 96-well plate (CELLSTAR plate, 96w, F, νClear, TC, PS, black, lid, Greiner) 24h before the start of the assay in 100µL of 5% DCC, 1% PS RPMI-1640 and were either stimulated with DMSO or 10nM R1881 for 24 hours. All cells were gently washed with PBS and LNCaP-eGFP cells were co-cultured in DMSO with either 3750 DMSO- or 3750 R1881 stimulated M1- or M2-polarized macrophages. Cell proliferation was measured with the IncuCyte Zoom fluorescent signal imaging system for 7 days. Data was normalized to time point zero (t = 24hrs) to account for possible fluorescence intensity artifacts upon initialization. To compare Incucyte results to *in silico* results, PCABM data was normalized to the number of tumor cells upon initialization.

Cell apoptosis was measured and analyzed using IncuCyte Zoom (EssenBioScience) on similar cell numbers, timespans and set-up as described previously with 0.5 mM Caspase-3/7 Red Reagent for Apoptosis (Essen BioScience), while apoptosis control was induced by supplementing to 0.5 mM Phenylarsine Oxide (PAO, Sigma). To investigate growth of LNCaP-abl cells in androgen-deprived conditions, 250 LNCaP-abl cells were seeded on a 96-well plate and cultured in RPMI-1640, 5% DCC + 1% PS. Cell proliferation was measured and analyzed by brightfield analysis with the IncuCyte Zoom (Essen BioScience) for 10 days.

#### Agent Based Model Design

Our two-dimensional PCABM consists of four agents (cell types): tumor cells, M1 and M2 polarized macrophages, and fibroblasts as these are the most abundant cell types and key players in the prostate TME^44^. PCABM requires specific size grid cells, although in reality actual cell sizes vary, therefore each grid cell was assigned the size of one tumor cell^45^ as 142.89 *µ*m^2^. Agents occupy exactly one position on a customizable rectangular grid, which size was scaled to *in vitro* well size leading to a 125x125 square grid (reality: 1.48 mm^2^).

To emulate *in vitro* settings, different agent types are randomly scattered on the grid upon initialization, with seeding densities matching *in vitro* experiments. PCABM runs for a fixed number of time steps of four hours every simulation, and each cell type has a probability to perform actions in the order: tumor cells, fibroblasts, M1 macrophages, and M2 macrophages (summarized in **Figure 1A**).

**Figure 1:**
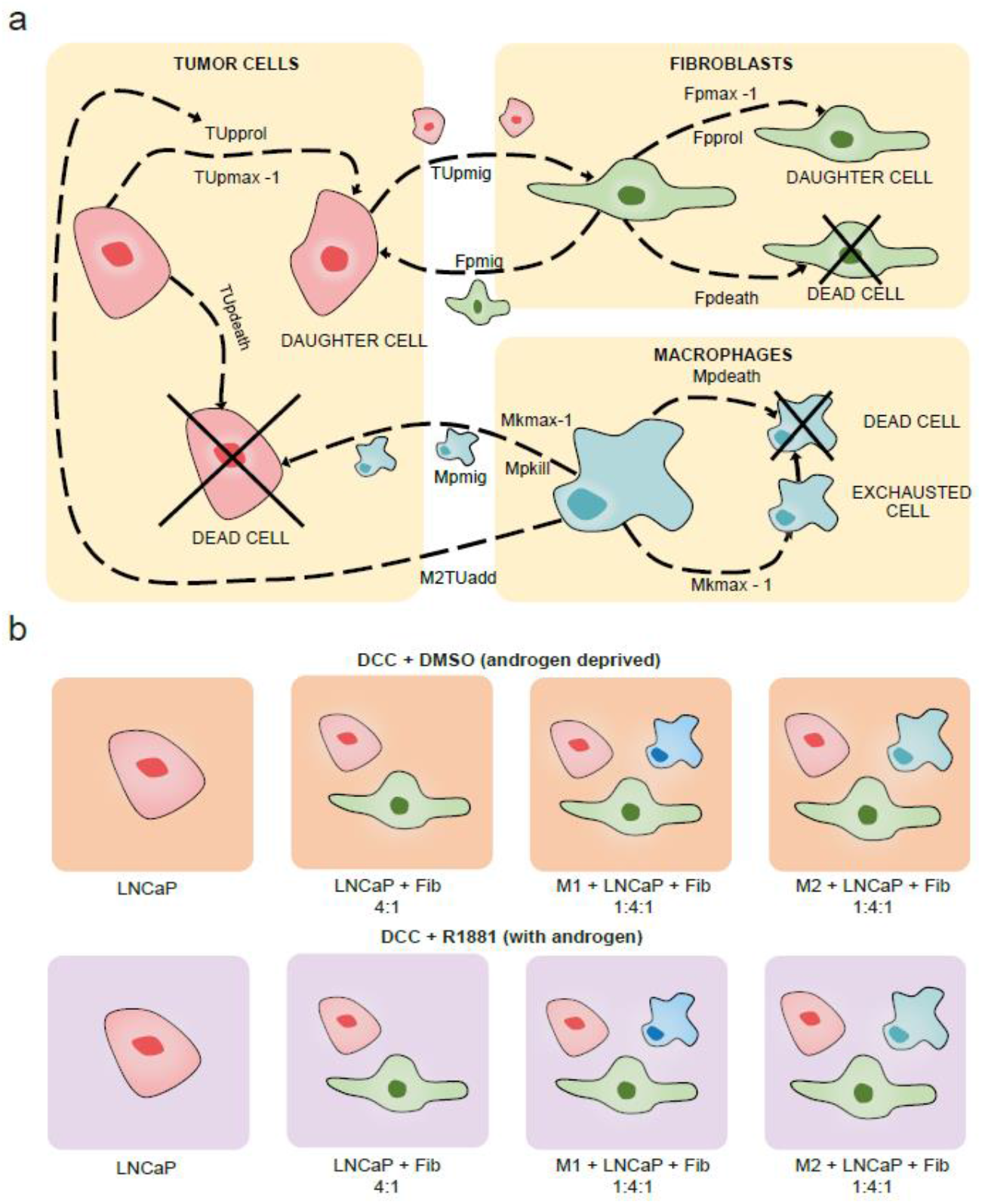
PCABM parameter and cell type action overview. **a)** Overview of all modelled cell interactions, in which each cell type can migrate, idle and die. Tumor cells and fibroblasts proliferate, while macrophages can either kill or support tumor cells depending on their subtype. **b)** PCABM is optimized for two in vitro co-culture conditions: cells grown in dextran coated charcoal (DCC) supplemented medium without androgen (DMSO, upper panels) and with androgen (R1881, lower panels). The different cell types are LNCaP, LNCaP + fibroblasts and LNCaP + fibroblasts + either M1 or M2-polarized macrophages.

Tumor cells can proliferate (TU_pprol_), die (TU_pdeath_, spontaneous death) or migrate (TU_pmig_) either towards fibroblasts or in random directions (TU_rwalk_ and have limited proliferation capacity (TU_pmax_). Fibroblasts can proliferate (F_pprol_) with limited capacity (F_pmax_), die (F_pdeath_, spontaneous) or migrate (F_pmig_) either towards tumor cells or in random directions (F_rwalk_). M1 and M2 polarized macrophages can migrate (M_pmig_) either towards tumor cells or randomly (M_rwalk_). Macrophages can kill (M_pkill_) when bordering a tumor cell, with maximum killing capacity (M_kmax_) before exhaustion and can spontaneously die (M_pdeath_). M2 polarized macrophages were calibrated to have attenuated tumoricidal activity compared to M1 polarized macrophages. Additionally, M2 polarized macrophages have the ability to increase tumor cell proliferation probability (M2_TUadd_).

Migration and proliferation processes requires unoccupied grid space in all agents’ neighborhood (Moore neighborhood), such that agents compete for space upon performing actions. Finally, inactive agents idle. All actions have calibrated stochastic probabilities, which resembles stochasticity observed in biological processes. An overview of model parameters is shown in **Supplementary Table S1**.

#### Initial parameter estimation

Tumor cell and macrophage migration parameters from Kather *et. al.*^40, 41^ were scaled to match PCABM grid size and time steps. Other parameter values were estimated using particle swarm optimization (PSO), which uses swarm behavior to search for global solutions^46^ and has been useful in a variety of optimization problems, including ABM^40, 47–49^.

Relative tumor cell numbers produced by PCABM were compared to *in vitro* relative growth curves to estimate parameters. TU_pmax_ was assumed to be the same in presence or absence of hormone and estimated only in hormone proficient conditions, in which ADT is assumed to be non-toxic. TU_pprol_ was instead fitted independently in the two hormonal conditions. The tumor cell apoptotic probability was measured *in vitro* using a caspase 3 and 7 assay and was assumed equal for both androgen pro- and deficient conditions. PSO was ran 50 times for each biological replicate (replicate optimizations in **Supplementary Figure S2**), with fixed parameter set to the median of the triplicate to be used as input for the next PSO iteration.

Similar to tumor cells, relative fibroblast numbers produced by PCABM were compared to relative fibroblast growth curves *in vitro* and parameters F_pprol_, F_pmax_ and F_pdeath_ were fitted. Fibroblast parameters were only optimized for DCC+R1881 conditions, since fibroblasts exhibit similar growth curves in androgen pro- and deficient conditions (**Supplementary Figure S2**)^21^. Fibroblast migration parameters (F_pmig_ and F_rwalk_) and tumor cell migration towards fibroblasts (TU_rwalk_) were qualitatively tuned by comparing model visualizations to *in vitro* captured cell dynamics.

Macrophage optimizations were performed separately for M1- and M2-polarized macrophages in the presence of both tumor and fibroblast cells for both DMSO and R1881 conditions (**Supplementary Figure S3-4**). Again, PCABM relative tumor cell numbers in macrophage presence were fitted to *in vitro* relative tumor cell numbers. The parameters M1_pkill_ and M1_kmax_ were optimized in hormone proficient conditions and killing capacities were assumed at maximum in these conditions as justified by our *in vitro* killing observations (**Supplementary Figure S5**). However, for vehicle conditions only M1_pkill_ was optimized, as this value is reasonably lower in hormone deficient conditions. Similarly, M2-polarized macrophage killing M2_pkill_ was optimized with M2_kmax_ the same as M1_kmax_, although simultaneously M2_TUadd_ was optimized as tumor promoting growth parameter (**Supplementary Figure S5**). A full list of the estimated parameters can be found in **Supplementary Table S2**.

#### Exploring effects of ADT on the prostate TME

Simulations solely included tumor cells and macrophages to exclude possible confounding effects of fibroblasts. Parameters were estimated similarly to previous parameter optimizations, optimizing 50 times with PSO in triplicate. However, instead of fixing the median parameter value over all triplicates to create one model, median parameters were fixed for each triplicate model individually. Killing probability (M_pkill_) and capacity (M_kmax_) of macrophages were estimated separately for M1- and M2-macrophages in hormone proficient conditions.

#### Modeling castration resistance

Using PCABMs optimized hormonal TME conditions, CRPC growth was simulated by seeding a co-culture of androgen sensitive and resistant tumor cells (1:100) in hormone deprived conditions. Resistant tumor cells have different proliferation probability and capacity parameters (TU_pprolres_ and TU_pmaxres_ respectively), which were fitted to *in vitro* growth of LNCaP-abl cells (androgen ablated), an ADT resistant clone derived from LNCaP cells. Resistant tumor cells migrate as fast as non-resistant cells and have the same probability of spontaneous death as non-resistant tumor cells in hormone proficient conditions. To simulate interactions in the TME upon CRPC development also fibroblasts, M1- or M2-macrophage agents were added. Since the amount of TME cell infiltration varies in prostate tumors, simulations were run with various ratios of different cell types.

#### Patient samples and histology

Spatial cellular patterns produced by PCABM were compared with a histological sample from a radical prostatectomy specimen, which was formalin fixed, paraffin embedded (FFPE). Tissue was stained with hematoxylin and eosin (HE) and a 200x enlarged microscopy image was taken.

#### Statistical analysis

Statistical analysis of growth rate differences in hormone conditions was performed using linear mixed-effect models with longitudinal analysis using R package *TumGrowth*^50^. For validation, *in vitro* LNCaP cell growth was tested in different hormone conditions over time and also PCABM output for CRPC simulations with different cell types was analyzed similarly. Different TME compositions were tested for effects on simulated relative tumor cell number over time. Type II analysis of covariance (ANOVA) with Wald tests were used to calculate p-values with significance cutoff 0.05.

#### Data and code availability

The model used in this study is publicly available in https://github.com/SysBioOncology/PCABM_ADT.

## Results

### PCABM conceptual model

We developed an ABM consisting of tumor cells, fibroblasts, M1 and M2 macrophages, which are seen as agents and scattered randomly on grid upon initialization to mimic *in vitro* settings. These cellular agents perform actions (proliferate, die) and interact with each other as schematically represented in **Figure 1A** (see **Material and Methods** for a more extensive description of the model).

We optimized the PCABM on co-cultures experimental data (six technical replicates spanning three biological replicates) measured in androgen proficient R1181 conditions versus hormone deprived vehicle control conditions to mimic the TME in normal and ADT conditions respectively (**Figure 1B**).

### PCABM forms similar growth patterns as in vitro co-cultures and histological samples

Upon initialization of PCABM, cells are randomly distributed across a grid and self-organized to form complex spatial patterns over time (**Figure 2A**). In our *in silico* PCABM, we observe similar spatial growth patterns to those observed *in vitro* (**Figure 2B**) and to those observed in human tumor samples, as identified in HE stained formalin-fixed paraffin embedded prostate tumor tissue (**Figure 2C**). These observations illustrate PCABM’s ability to reliably model spatial PCa growth pattern complexity *in silico* from a simple set of assumptions and optimizations.

**Figure 2:**
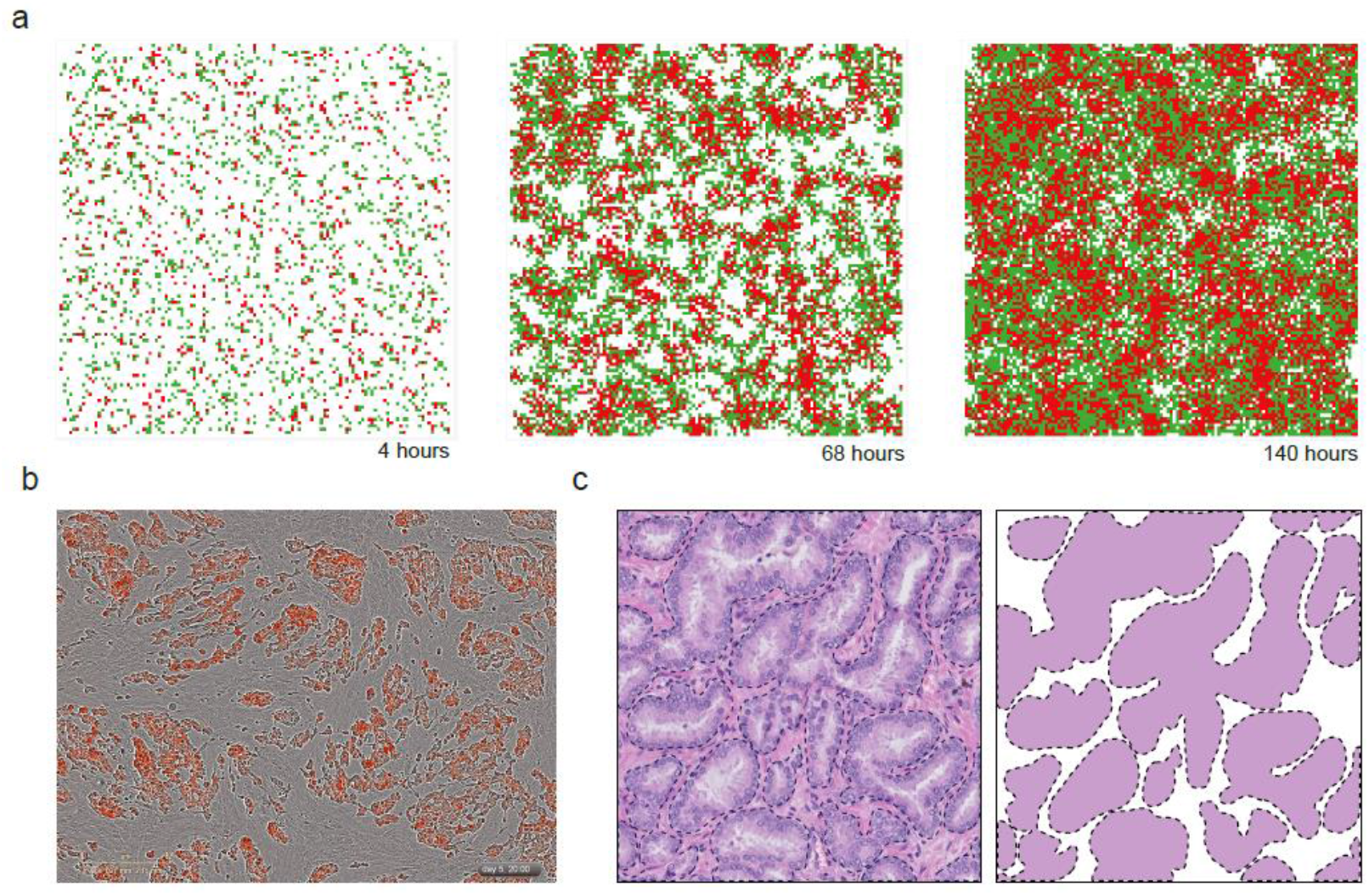
Prostate TME spatial patterns in silico, in vitro, and in vivo in hormone proficient conditions. **a)** Modeled tumor cells (red) and fibroblasts (green) are randomly distributed across PCABM lattice, but spatiotemporally organize after 4, 68 and 140 hours of pseudo-time. **b)** In vitro co-culture of tumor cells (red) and fibroblasts (brightfield, 1:1 ratio) after 140 hours. **c)** FFPE HE staining at 200x magnification of a primary prostate tumor, showing distinct epithelial tumor foci (masked image) surrounded by stroma.

### Hormonal response of PCa cells is accurately captured by PCABM

PCABM simulations recapitulate LNCaP cell growth curves observed in *in vitro* experiments well in both hormone proficient and deficient conditions (**Figure 3**). Model estimation of tumor cell proliferation (TU_pprol_) shows a threefold increase in tumor cell proliferation as response to R1881 treatment (TU_pprol_ = 0.1144 for R1881 versus 0.0389 vehicle control; **Supplementary Figure S2** for parameter optimizations). When adding fibroblasts *in silico* to the culture under R1881 conditions, a slight reduction in in the growth rate is observed without changing proliferation parameters, matching the corresponding experimental data (**Figure 3**). This change underlines the predictive power for ABM stochastic modeling without additional adjustments.

**Figure 3:**
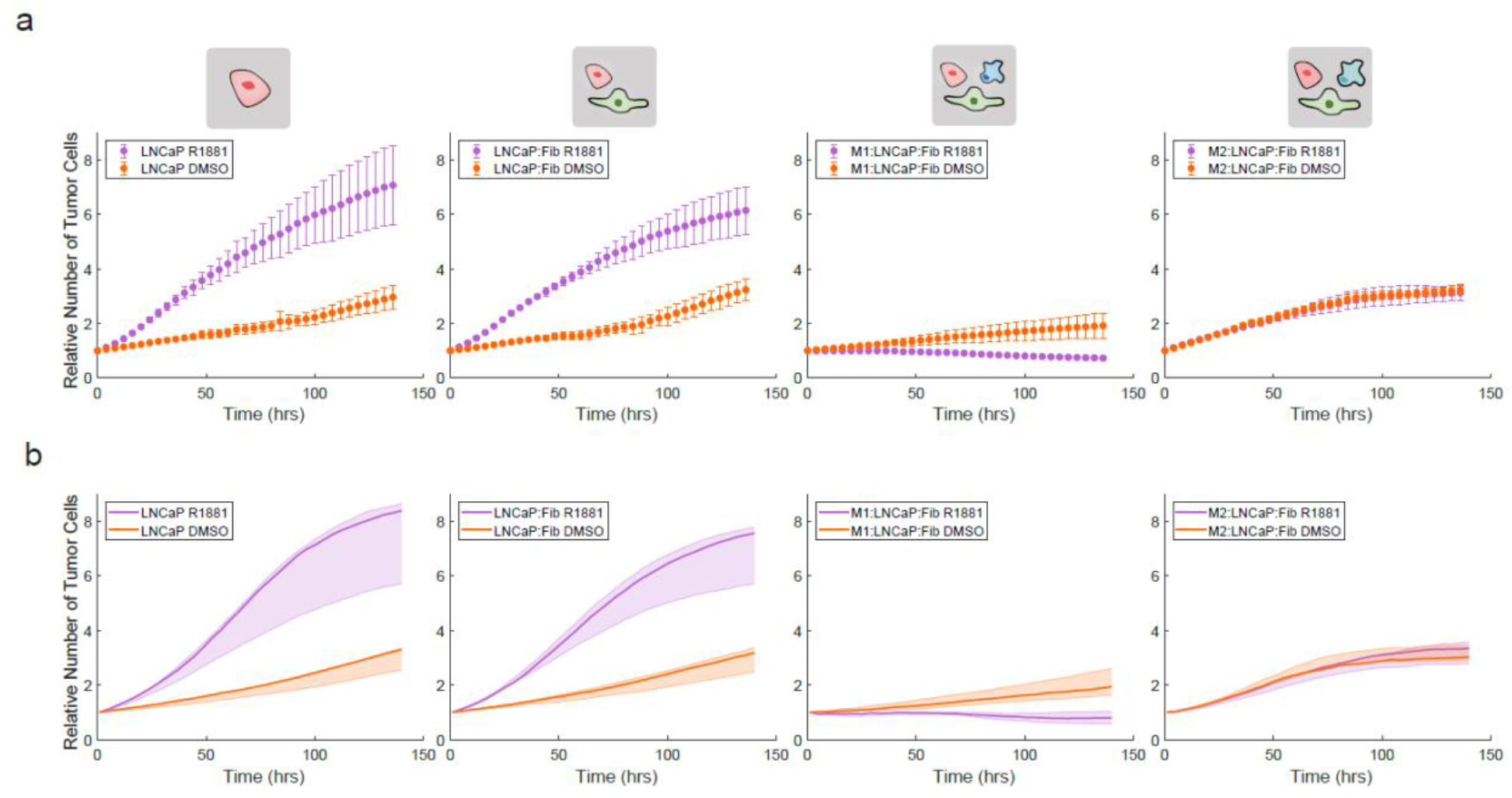
In vitro tumor cell proliferation and hormone response is accurately captured by PCABM’s optimized in silico parameters. **a)** Incucyte data for different co-cultures in hormone deficient (DMSO, orange) and hormone proficient (R1881, purple) conditions for sequentially LNCaP monoculture; LNCaP and fibroblast co-culture; LNCaP, fibroblast and M1-polarized macrophage co-culture; LNCaP, fibroblast and M2-polarized macrophage co-culture. **b)** PCABM model behavior after parameter optimization in hormone deficient (DMSO, orange) and hormone proficient (R1881, purple) in silico conditions for sequentially LNCaP monoculture; LNCaP and fibroblast co-culture; LNCaP, fibroblast and M1-polarized macrophage co-culture; LNCaP, fibroblast and M2-polarized macrophage co-culture. Data represent the average of three biological replicates, with six technical replicates each. Error bars indicate standard deviation. Lines represent PCABM model output with the median of optimized parameters over three biological replicates. Shading represents model output for optimized parameters within interquartile range given by 50 optimizations for each biological replicate.

Co-culturing M1-polarized macrophages together with LNCaP and fibroblast, we observed an *in vitro* strong decrease in tumor growth rate compared to LNCaP mono-cultures and LNCaP + fibroblast co-cultures, while such an effect was less apparent in the hormone deprived condition (**Figure 3A**). By simulating the same experimental condition (i.e. model with LNCaP, fibroblasts and M1 macrophages) and optimizing PCABM’s M1 macrophage killing probability (M1_pkill_) based on these data, we found a 22-fold decrease in killing capacity in hormone deficient (DCC+DMSO) versus hormone proficient (DCC+R1881) conditions (M1_pkill_ = 0.005 and 0.1116 respectively; **Figure 3B**). In contrast, replacing M1-like for M2-like polarized macrophages did not result in a differential effect in growth curves between hormone conditions both *in vitro* and *in silico* (**Figure 3A,B**).

Such cell culture growth dynamics could be reliably reproduced *in silico* using PCABM, with different observed tumor cell proliferation and kill capacities in the hormonal conditions for M2-polarized macrophages (TU_pprol_ R1881= 0.0389 and TU_pprol_ DMSO = 0.1348; M2_pkill_ R1881 = 0.0223, M2_pkill_ DMSO = 0.0348; **Figure 3B, Supplementary Figure S3-4**). Taken together, these data suggest that PCABM accurately describes PCa cell proliferation potential and the impact of R1881 treatment thereon, when co-cultured with different TME cell types.

### PCABM predicts immunomodulatory effects of ADT on macrophages

Through PCABM parameter optimization we further estimated whether the hormone-driven decrease of LNCaP cell growth in co-culture with M1 or M2 polarized macrophages was tumor cell intrinsic or related to macrophage tumoricidal activity. For this purpose, we cultured LNCaPs with macrophages but without the presence of fibroblasts and saw differences compared to previous growth rates, with a clear tumoricidal effect for M1 macrophages supplemented with R1881 (**Figure 4A**). Paradoxically, optimizing LNCaP TU_pprol_ in vehicle conditions while using macrophage M1_pkill_ and M1_kmax_ that we previously optimized in hormone-proficient conditions, resulted in higher predicted proliferation values (TU_pprol_ DMSO = 0.1550; TU_pprol_ R1881 = 0.1144, **Figure 4B, Supplementary Figure S4**). Since higher LNCaP TU_pprol_ is expected upon R1881 treatment, we optimized M1_pkill_ while keeping LNCaP proliferation constant on vehicle conditions (TU_pprol_ DMSO = 0.0389), which resulted in an improved PCABM fit to *in vitro* data with smaller mean square error (MSE) between data and model fit for all three *in vitro* replicates (**Figure 4B, Supplementary Figure S5**). Importantly, R1881 conditions increased M1_pkill_ capacity 21-46 fold (M1_pkill_ DMSO = 0.005 in vehicle control; M1_pkill_ R1881 = 0.2034). These PCABM optimizations suggest that changes in tumor cell viability upon hormone deprivation are not solely dictated by decreased tumor cell proliferation but are also impacted by M1 macrophage tumoricidal effects.

**Figure 4:**
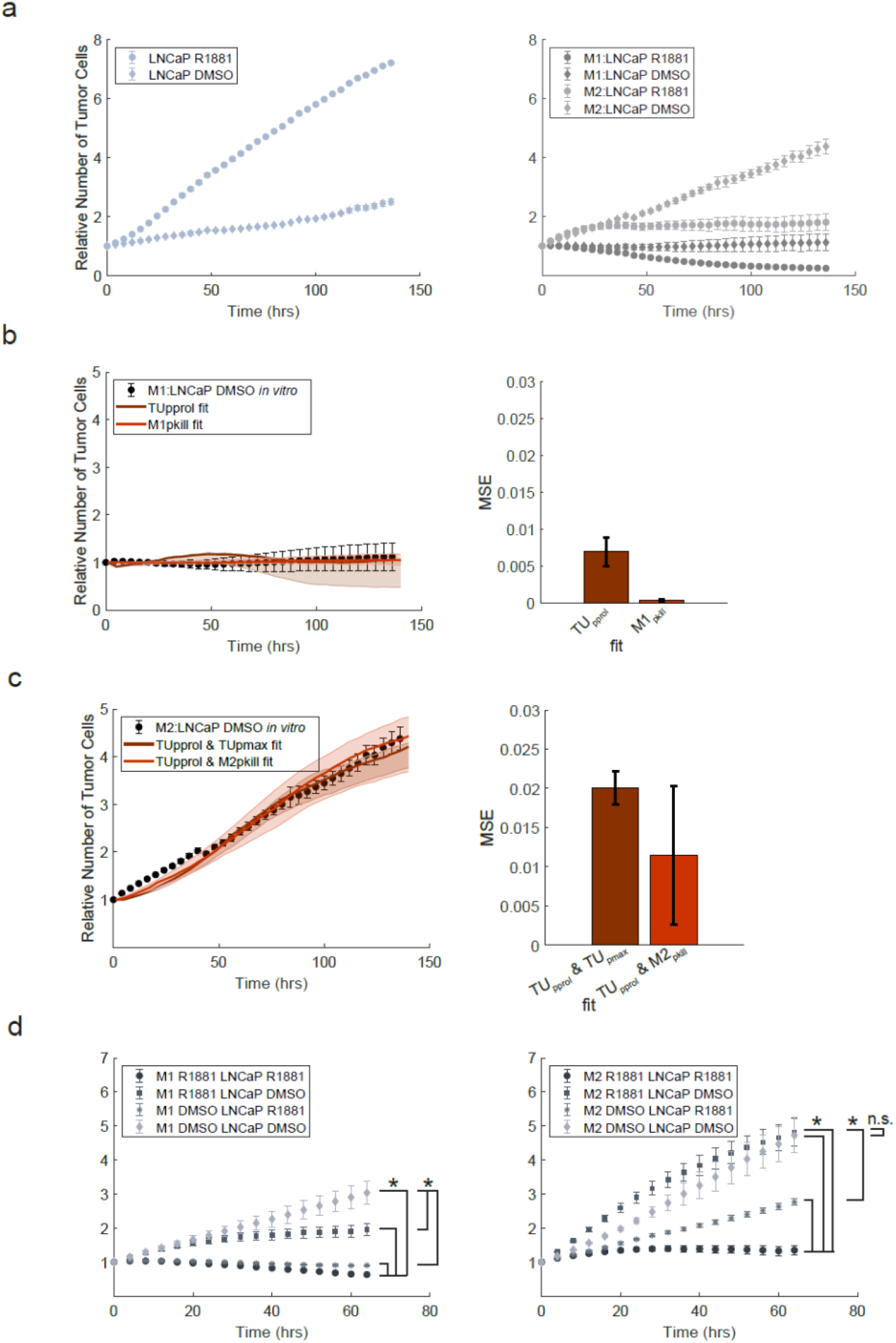
PCABM predicts immunomodulatory ADT-mediated macrophage tumoricidal effects. **a)** LNCaP growth curve alone (left) or in co-culture with M1- or M2-macrophages (right) in absence or presence of R1881. **b)** PCABM optimization for TU_pprol_ and M1_pkill_ in DCC+DMSO (left) and mean squared error (MSE) between experimental data and PCABM for M1:LNCaP TU_pprol_ and M1_pkill_ (right). **c)** PCABM optimization for TU_pprol_ and or M2_pkill_ in DCC DMSO (left) and mean squared error (MSE) between experimental data and PCABM for M2:LNCaP TU_pprol_ + TU_pmax_ and TU_pprol_ + M2_pkill_ (right). **d)** Growth curve of LNCaP co-cultured with M1-macrophages individually stimulated with DMSO or R1881 (left) and growth curve of LNCaP co-cultured with M2-macrophages individually stimulated with DMSO or R1881 (right). Bars and error represent mean and standard deviation over MSE of 50 optimizations for replicate 1. Dots represent average and error bars represent standard deviation of six technical replicates. Lines represent PCABM output with the median of optimized parameters. Shading represents model output for optimized parameters within interquartile range given by 50 optimizations.

To observe whether such an approach would also improve MSEs in the M2-polarized PCABM, and whether M2-macrophage polarization has differential effects on the TME compared to M1-polarized macrophages, we again optimized M2_pkill_ while keeping LNCaP proliferation constant to vehicle conditions (TU_pprol_ DMSO = 0.1341), with TU_pmax_ DMSO = 5, which only slightly improved PCABM fit and MSEs (**Figure 4C, Supplementary Figure S5**). As expected, PCABM indicates that M2-macrophages exhibit less tumoricidal activity compared to M1-macrophages and become tumor promoting in vehicle conditions, enhancing predicted tumor growth (TU_pprol_ 2-3 fold increase) while decreasing tumor killing capacity (M2_pkill_ 2-4 decrease) relative to R1881 conditions (TU_pprol_ DMSO = 0.0384 and TU_pprol_ R1881 = 0.1128 and M2_pkill_ DMSO = 0.0219 and M2_pkill_ R1881 = 0.0441; **Figure 4C, Supplementary Table S2**). In co-cultures, we validated these findings with individually stimulated co-culture cell constituents. For M1 co-cultures we observed that growth is significantly increased in hormone deprived conditions, while for M2 co-cultures this effect is not present (**Figure 4D**). These results suggest that ADT exerts an immunomodulatory effect on tumor cell killing.

### Spatial effects in the TME and differential macrophage tumoricidal capacities enhance TME cellular dynamics

We next sought to investigate how the TME contributes to the emergence of CRPC. Experimental data from castration resistant LNCaP-abl (androgen ablated) cells grown in hormone deprived conditions was used to fit proliferation parameters for resistant cells (**Materials and Methods, Supplementary Figure S2**). In contrast to LNCaP cells, *in vitro* LNCaP-abl growth increases exponentially in hormone deprived conditions (**Supplementary Fig. S2C**). Therefore, to mimic LNCaP-abl growth observed *in vitro*, we optimized a higher tumor cell proliferation (TU_pprolres_ = 0.06) for resistant cells, which is almost twice that of LNCaP TU_pprol_ in hormone deprived conditions. Interestingly, LNCaP-abl cells readily form clusters of resistant cells *in silico* (**Figure 5A**), which is also observed when growing LNCaP-abl cells *in vitro*.

**Figure 5:**
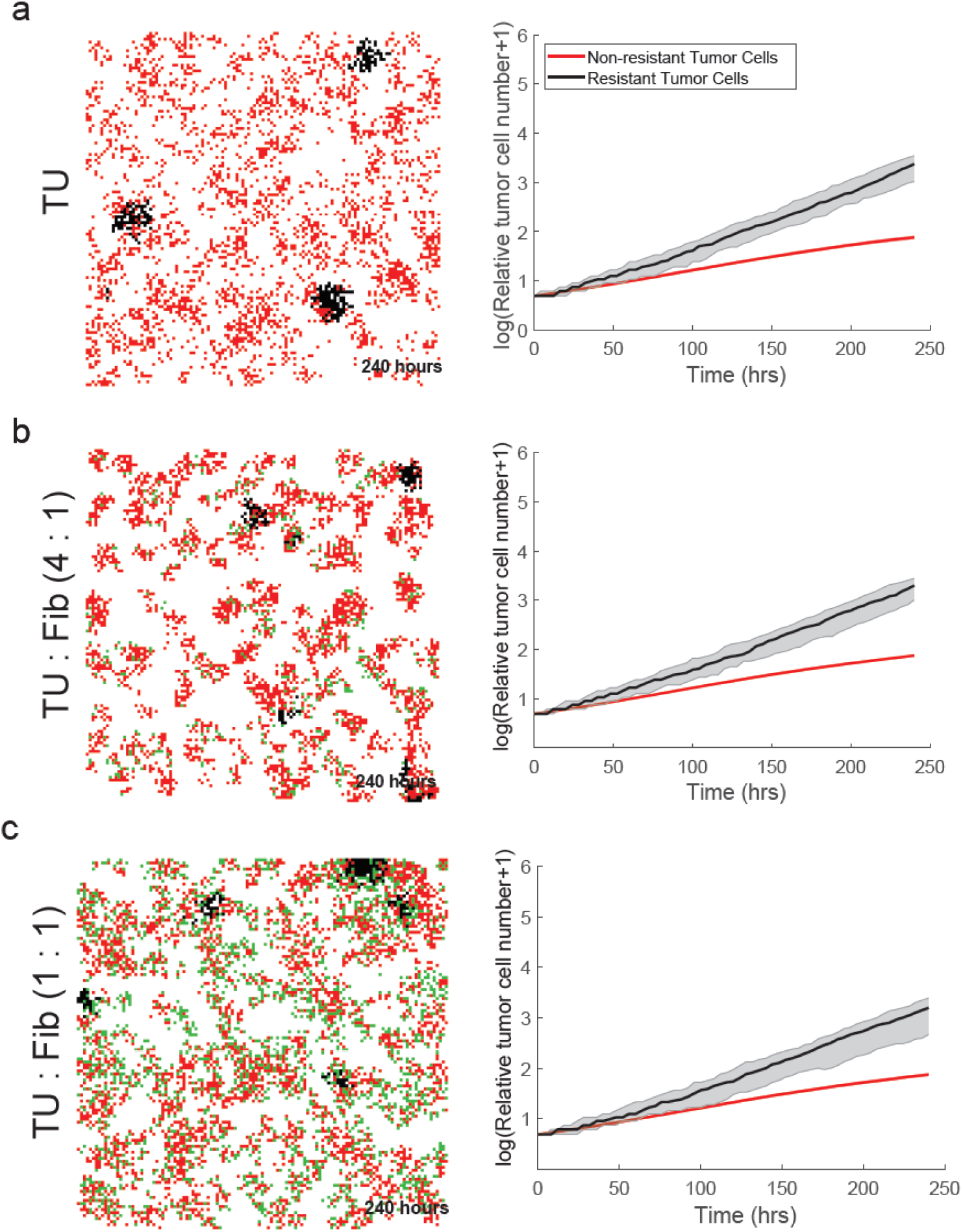
CRPC simulations in PCABM with fibroblasts. **a)** Growth of LNCaP and LNCaP-abl cells. **b)** Relative growth of tumor cells seeded with fibroblasts at a 4:1 ratio **c)** Relative growth of tumor cells seeded with fibroblasts at a 1:1 ratio For all panels, PCABM (left) is compared to Incucyte (right) of co-cultures of LNCaP cells with LNCaP-abl cells and fibroblasts.

While the *in silico* addition of fibroblasts does not affect proliferation speed, there are increased fibroblast directional migration effects towards tumor cells. These effects result in increased hormone-sensitive tumor-cell cluster formation, which in turn is balanced by cellular competition for space as fibroblasts take up growth space (**Figure 5B-C**). These data suggest that not only the population growth of TME constituents, but that also the available TME space is an important characteristic to describe the entirety of TME cellular growth dynamics.

### Macrophage phenotype and influx play a critical role on resistant tumor cell growth

Next, we further enriched our *in silico* model, by including tumoricidal M1 polarized macrophages in CRPC-PCABM, which has a repressing effect on both CRPC and hormone responsive PCa proliferation speed. Since the number of tumor-resident macrophages vary greatly between PCa samples^23, 51^, which can be partially explained by differences in tumor volume and macrophage influx, we wondered how PCABM would respond to varying levels of macrophages. When quadrupling the amount of macrophages to tumor cells, tumor cell population extinction is quickly achieved *in silico* (**Figure 6A,B**). Interestingly, the addition of a large fibroblast presence seems to reduce macrophage tumoricidal effects (**Figure 6B**). Conversely, M2-polarized macrophages significantly increase tumor cell proliferation, and proportionally to a larger extent for CRPC as compared to hormone-sensitive PCa cells (**Figure 6D,E**). Additionally, when changing the ratios between tumor cells and M2-polarized macrophages we observe a growth reduction of both resistant and hormone-sensitive tumor cells (**Figure 6D,E**). Taken together, these observations demonstrate how a higher influx of macrophages lead to tumor remission even in the context of resistant tumor cells, while fibroblasts provide a protective niche for resistant tumor cells to proliferate in.

**Figure 6:**
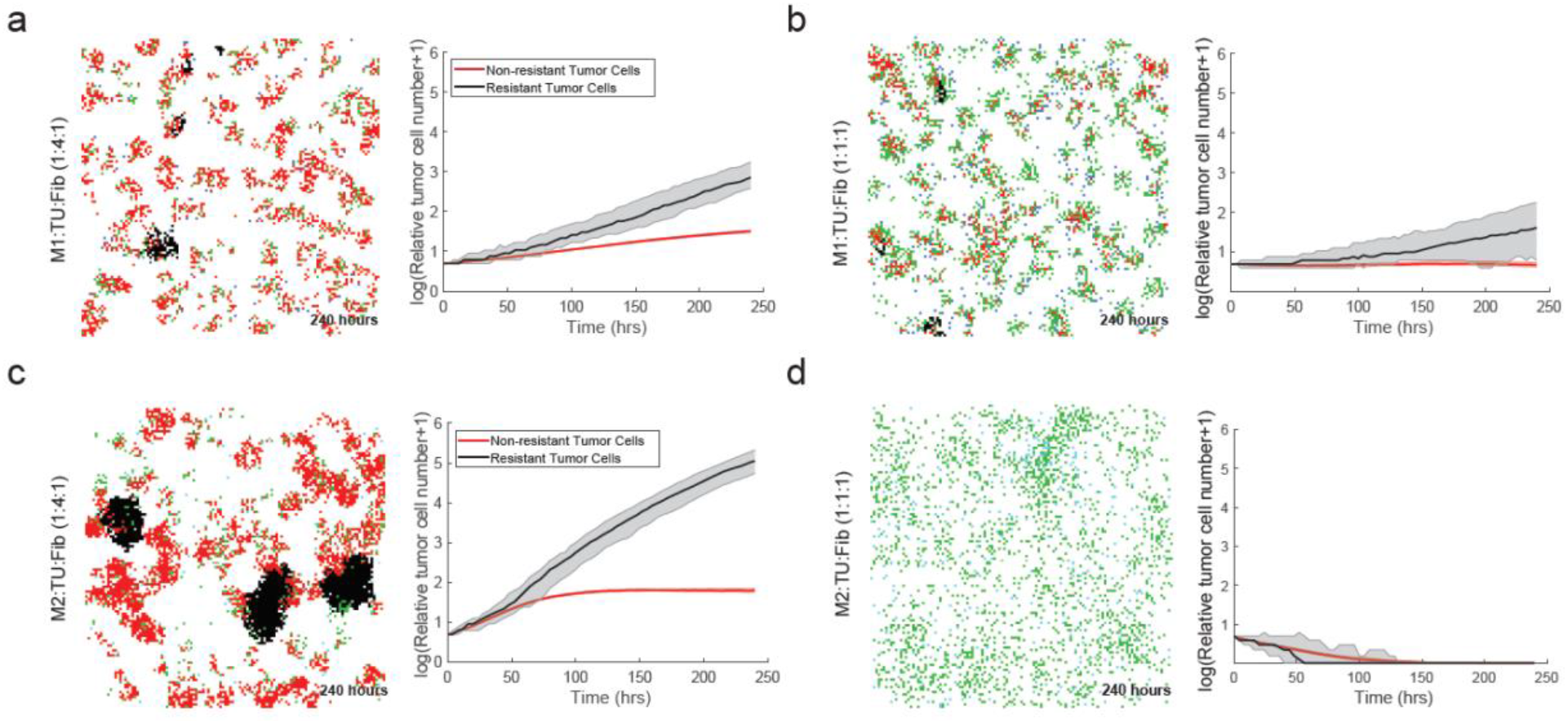
CRPC simulations in PCABM with fibroblasts and either M1 and M2 polarized macrophages. **a)** M1 macrophages seeded with tumor cells and fibroblasts at a 1:4:1 ratio **b)** M1 macrophages seeded with tumor cells and fibroblasts at a 1:1:1 ratio **c)** M2 macrophages seeded with tumor cells and fibroblasts at a1:4:1 ratio **d)** M2 macrophages seeded with tumor cells and fibroblasts at a 1:1:1 ratio For all panels, PCABM (left) is compared to Incucyte (right) of co-cultures of LNCaP cells and LNCaP-abl cells, fibroblasts and M1- or M2-polarized macrophages.

## Discussion

Because AR plays a key role in PCa progression, patients with metastatic disease recurrence are typically treated with AR-targeted therapeutics^52, 53^. Since cells in the TME also express AR, they are consequently also affected by ADT, which could affect cell-cell interactions. In this work, we replicated ADT-conditions *in silico* in a PCa-specific ABM, which is able to model the spatiotemporal complexity of prostate TME cell interactions in both hormone pro- and deficient conditions. By implementing a simple set of stochastic assumptions, an intrinsically organized, self-assembling TME cellular structure emerges in PCABM that resembles the histology in PCa patient samples. Since PCa is multifocal in 60-90% of cases^54^, these simulated tumor foci further underscore the ability of the PCABM to form clinically relevant spatial patterns and suggest that the TME plays a critical role in the formation of multifocal disease.

Our modeling assumptions were calibrated and refined using data from extensive *in vitro* co-cultures, that incorporate cell proliferation and migration data. Because PCABM is currently modeled only for LNCaP cells, it is currently limited in its ability to accurately replicate PCa growth and development of CRPC. However, the model is adaptable to other AR-positive PCa cell lines, provided that *in vitro* data exists for calibration demonstrating its strength in that the parameters are easily adaptable to other hypotheses. Multiple PCa cell lines have been developed with a wide variety of proliferation kinetics and response to hormones, which may lead to different PCABM results.

Recently, we found that AR plays a key-role in macrophage-mediated killing, being a critical tumor-intrinsic regulator and preventing macrophages from killing tumor cells in androgen deprived conditions^43^. Fully in line with this, our PCABM predicts that ADT affects the cellular behavior of both tumor cells and M1 macrophages, further solidifying our observation that AR plays an immunomodulatory role in the prostate TME. Independent *in vitro* experiments validated this, suggesting that ADT affects the differentiation of this cell type, which may potentially stimulate tumor growth. Interestingly, the addition of fibroblasts to

PCABM stimulates directional migration of both tumor cells and fibroblasts, resulting in a limited amount of space around the tumor cells. In androgen proficient conditions such a proliferation space will be severely limited due to high proliferation rates, whereas in androgen deficient conditions, such an effect will theoretically be less pronounced due to decreased proliferation rates of AR-responsive cells. These results suggest that fibroblasts block the access of M1 macrophages to tumor cells by their preferential clustering around tumor cells. Since macrophages are able to kill tumor cells through cell-to-cell contact^40^, fibroblasts may prevent macrophages from completing their tumoricidal activity.

In addition, we modeled CRPC formation in PCABM and showed that resistant cells form separate clusters due to the directional migration effects of fibroblasts. These findings support the multifocality of PCa and further highlight the tumor-protective role of fibroblasts by limiting the physical access of macrophages while creating a niche for tumor cells. Previously, the amount of stroma has been shown to be inversely correlated with recurrence-free survival, suggesting that stromal cells may protect tumor cells from being killed^55, 56^. Supporting this, M1 macrophages decreased the growth of both androgen-sensitive and -insensitive PCa cells, whereas M2 macrophages allowed castration-resistant tumor cells to rapidly take over the TME. Recently, tumor-associated macrophages have been associated with PCa progression after ADT^12^ and the development of CRPC^8^, which is supported by our findings on the immunomodulatory effects of ADT and CRPC growth. These findings are also consistent with our recent report, in which we showed that AR signaling in macrophages plays a critical role in PCa migration and invasion through TREM-1 signaling and a concomitant upregulation of IL-10^29^. In contrast, when AR signaling is blocked in CAFs, PCa cells migrate under the influence of upregulated CCL2 and CXCL8 secretion^25^. These studies further underline the tumor-driving effects of the prostate TME induced by ADT, along with the differential intercellular interactions in this context.

Technically, PCABM has been calibrated to *in vitro* time scales and data. For more *in vivo*-like PCABM representations, longer pseudo timescales are needed, and the currently modeled pseudo timescales could be extended with long-term culture data, although long-term culture has practical limitations. We approximated the prostate TME by including tumor cells, fibroblasts and macrophages, which are the most abundant cell types in PCa^44^.

In conclusion, we present PCABM, an *in silico* tool that simulates and accurately describes the functional interplay between prostate TME cells in hormone proficient and ADT conditions and in the emergence of CRPC. Our findings suggest that targeting TME cell types may provide a novel avenue for the treatment of CRPC, as different TME cell types influence castration-resistant tumor cell growth. In future research, PCABM could be used to design targeting strategies involving the TME to achieve optimal anti-tumor efficacy, which may serve as a blueprint for implementation in other cancer types.

## Acknowledgements

We express gratitude to all members of the Zwart and Bergman lab, and members of the NKI Oncogenomics division for helpful scientific discussion. We would like to thank Margot Passier for testing the code. This work was supported by Prostate Cancer Foundation, Department of Defense, Oncode Institute and Alpe d’HuZes/ KWF Dutch Cancer Society.

## Author contributions

A.Z., J.K., A.M.B, W.Z. and F.E. conceived the study. M.v.G. performed all *in silico* modeling experiments. A.Z. and J.K. carried out *in vitro* experiments. E.B. provided histology samples. M.v.G., A.Z., J.K., A.M.B. and W.Z. wrote the manuscript with input from all authors. A.M.B., W.Z. and F.E. supervised the study.

