## Supplementary Figures S1-6 for "Agent-based modeling of the prostate tumor microenvironment uncovers spatial tumor growth constraints and immunomodulatory properties"

Supplementary Figures S1-6

### Supplementary Figure 1

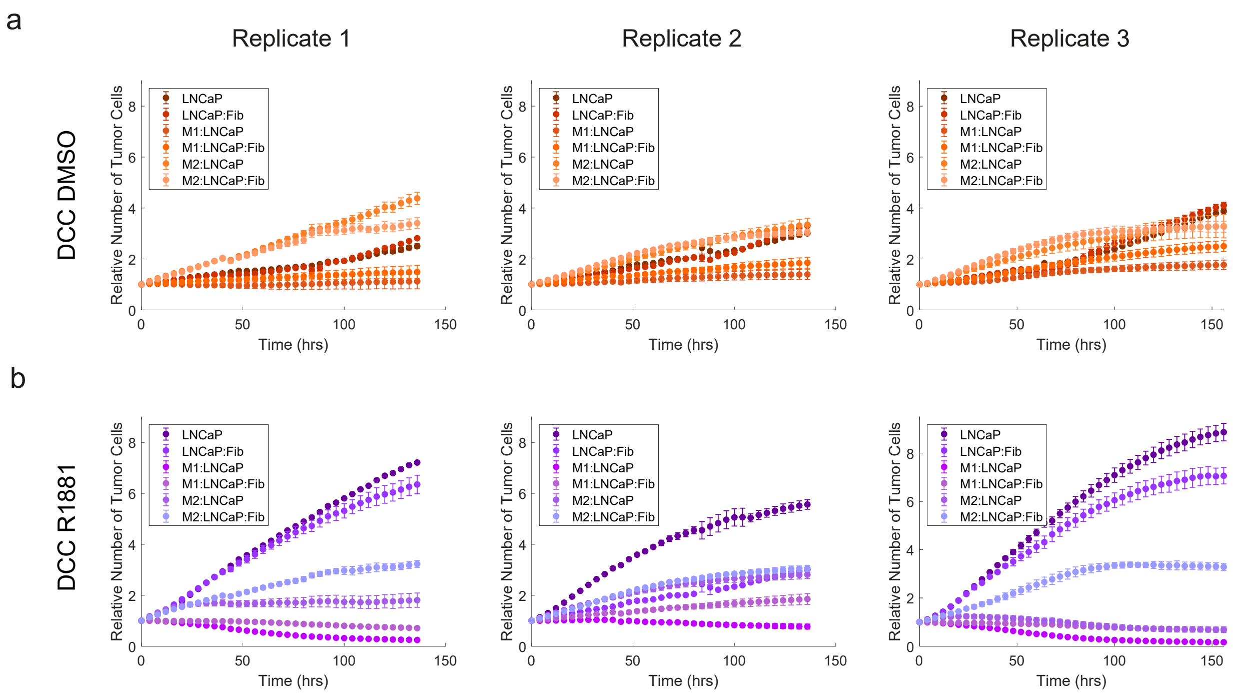

**Supplementary Figure 1: Biological replicates of LNCaP co-cultures in hormone deficient and proficient conditions.**

**a)** Three biological replicates of LNCaP cells co-cultured with fibroblasts alone, M1-or M2-macrophages alone or co-cultured with both cell types in hormone deprived conditions.

**b)** Three biological replicates of LNCaP cells co-cultured with fibroblasts alone, M1-or M2-macrophages alone or co-cultured with both cell types in hormone proficient conditions.

Dots represent the average of six technical replicates. Error bars represent the standard deviation.

### Supplementary Figure 2

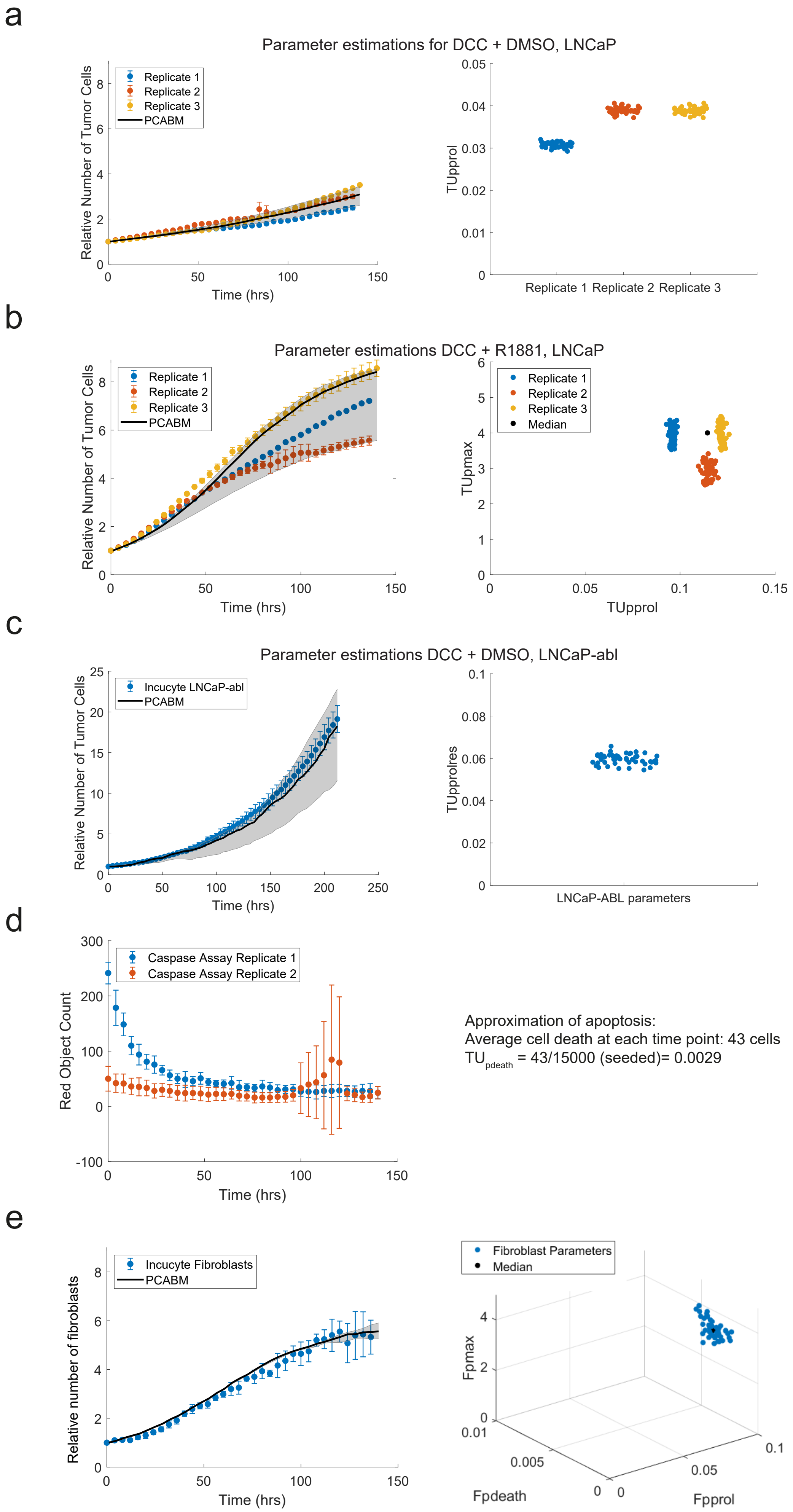

**Supplementary Figure 2: Particle Swarm Optimization estimations of parameters for LNCaP cell growth, apoptosis and LNCaP cell growth co-cultured with fibroblasts.**

**a)** Three biological replicates of LNCaP growth curves in hormone deficient conditions (left, dots represent Incucyte data, solid line represents PCABM). Estimation of the parameter $TU_{pprol}$ in hormone deficient conditions using three biological replicates (right).

**b)** Three biological replicates of LNCaP growth curves in hormone proficient conditions (left, dots represent Incucyte data, solid line represents PCABM). Estimation of parameters $TU_{pprol}$ and $TU_{pmax}$ in hormone proficient conditions using three biological replicates (right).

**c)** LNCaP-abl growth data (Incucyte) overlain with PCABM output with optimized parameters (left). Estimation of TUpprolres (right).

**d)** Incucyte red object count for caspase 3/7 assay with LNCaP cells. An object is interpreted as a dying cell.

**e)** Fibroblast growth data (Incucyte) projected on PCABM output with optimized parameters (left). Estimation of $F_{pmax}$, $F_{pdeath}$ and $F_{pprol}$ (right).

Optimized parameter sets (50 optimizations) for each biological replicate. For Incucyte data, dots represent the mean and error bars represent the standard deviation of six technical replicates. For PCABM data, lines represent *in silico* model output with median optimized parameters. Shading represents model output for optimized parameters within interquartile range for 50 optimizations per biological replicate.

### Supplementary Figure 3

a

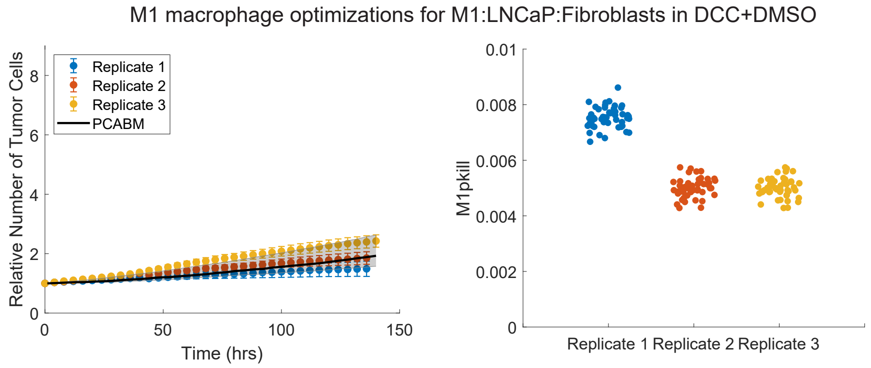


b

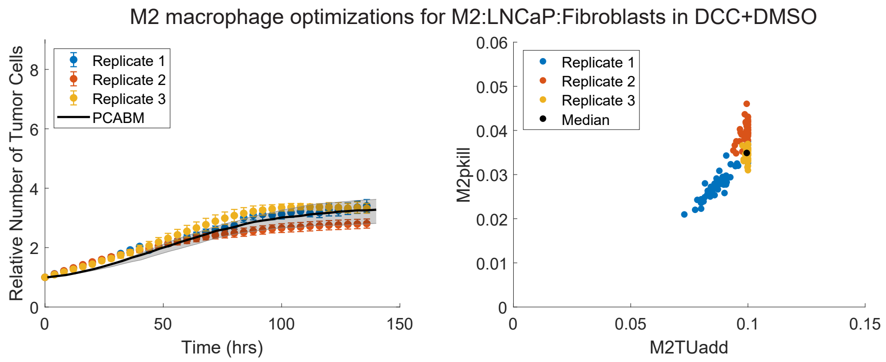

**Supplementary Figure 3: PCABM parameter estimations for LNCaP cells co-cultured with fibroblasts and M1-or M2-polarized macrophages in hormone deficient conditions.**

**a)** Three biological replicates of growth curves of LNCaP cells co-cultured with fibroblasts and M1 macrophages is hormone deficient medium (left, dots represent Incucyte data, solid line represents PCABM). Estimation of $M1_{pkill}$ in hormone deficient conditions (right).

**b)** Three biological replicates of growth curves of LNCaP cells co-cultured with fibroblasts and M2 macrophages is hormone proficient medium (left, dots represent Incucyte data, solid line represents PCABM). Estimations of $M2_{pkill}$ and $M2_{TUadd}$ in hormone deficient conditions (right).

Optimized parameter sets (50 optimizations) for each biological replicate.

For Incucyte data, dots represent the mean and error bars represent the standard deviation of six technical replicates. For PCABM data, lines represent *in silico* model output with median optimized parameters. Shading represent model output for optimized parameters within interquartile range for 50 optimizations per biological replicate.

### Supplementary Figure 4

a

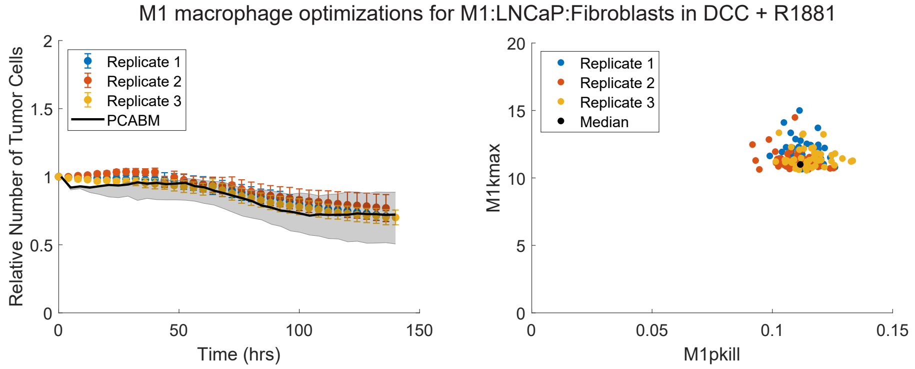


b

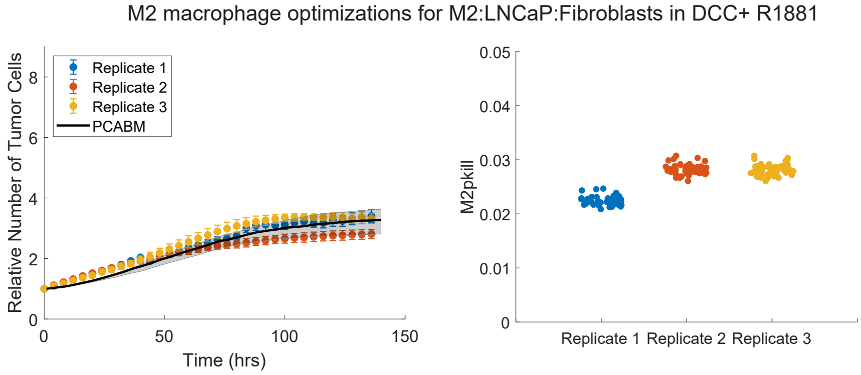

**Supplementary Figure 4: PCABM parameter estimations for LNCaP cells co-cultured with fibroblast and M1-or M2-polarized macrophages in hormone proficient conditions.**

**a)** Three biological replicates of growth curves of LNCaP cells co-cultured with fibroblasts and M1 macrophages is hormone proficient medium (left, dots represent Incucyte data, solid line represents PCABM). Estimations of $M1_{pkil}$l and $M1_{kmax}$ in hormone proficient conditions (right).

**b)** Three biological replicates of growth curves of LNCaP cells co-cultured with fibroblasts and M2 macrophages in hormone proficient medium (left, dots represent Incucyte data, solid line represents PCABM). Estimations of $M1_{pkill}$ and $M1_{kmax}$ in hormone proficient conditions (right). Optimized parameter sets (50 optimizations) for each biological replicate. For Incucyte data, dots represent the mean and error bars represent the standard deviation of six technical replicates. For PCABM data, lines represent *in silico* model output with median optimized parameters. Shading represent model output for optimized parameters within interquartile range for 50 optimizations per biological replicate.

### Supplementary Figure 5

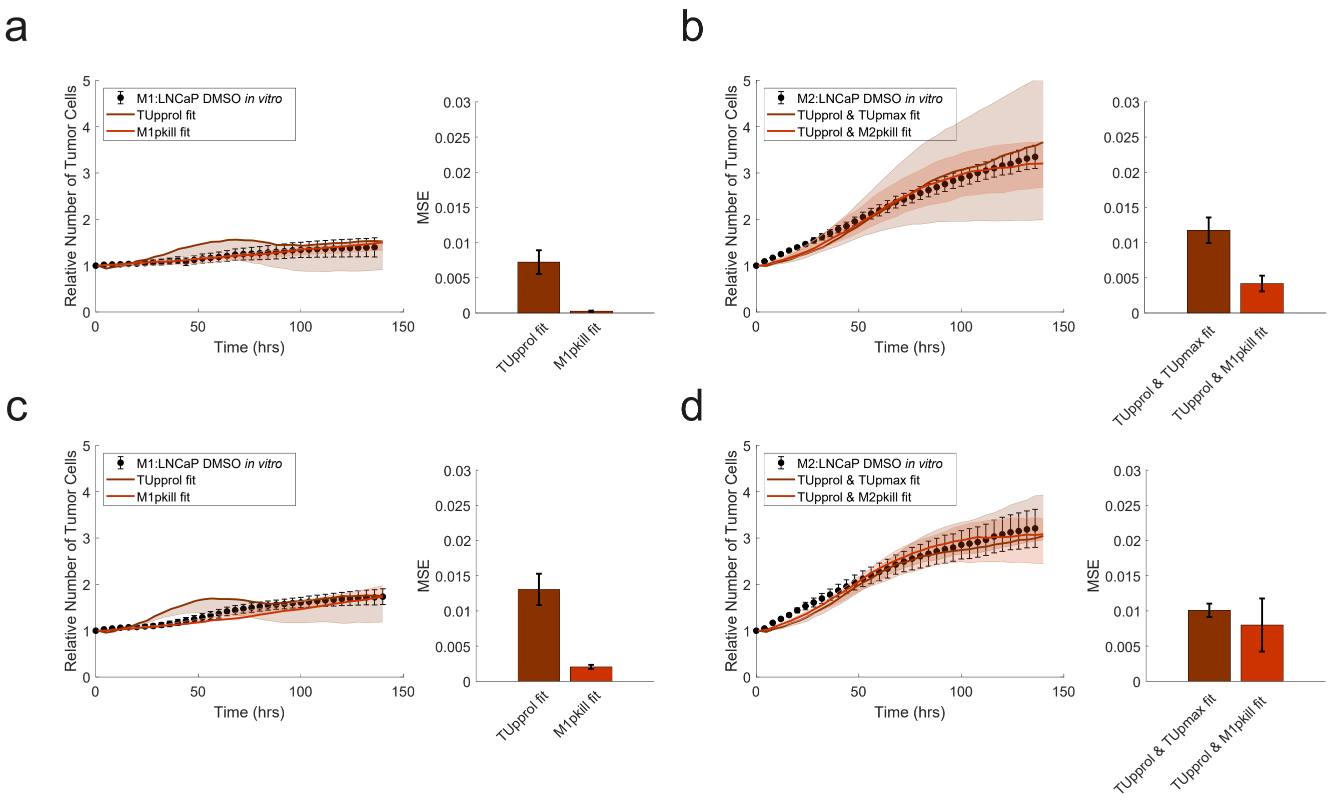

**Supplementary Figure 5: PCABM optimization for macrophage conditions.**

**a)** Optimization replicate 2 M1:LNCaP. PCABM optimization for $TU_{pprol}$ and $M1_{pkill}$ in DCC+DMSO (left) and mean squared error (MSE) between data and PCABM for M1:LNCaP $TU_{pprol}$ and $M1_{pkill}$ (right).

**b)** Optimization replicate 2 M2:LNCaP. PCABM optimization for $TU_{pprol}$ + $TU_{pmax}$ and $TU_{pprol}$ + $M2_{pkill}$ in DCC DMSO (left) and mean squared error (MSE) between data and PCABM for M2:LNCaP $TU_{pprol}$ + $TU_{pmax}$ and $TU_{pprol}$ + $M2_{pkill}$ (right).

**c)** Optimization replicate 2 M1:LNCaP. PCABM optimization for $TU_{pprol}$ and $M1_{pkill}$ in DCC+DMSO (left) and mean squared error (MSE) between data and PCABM for M1:LNCaP $TU_{pprol}$ and $M1_{pkill}$ (right).

**d)** Optimization replicate 2 M2:LNCaP. PCABM optimization for $TU_{pprol}$ + $TU_{pmax}$ and $TU_{pprol}$ + $M2_{pkill}$ in DCC DMSO (left) and mean squared error (MSE) between data and PCABM for M2:LNCaP $TU_{pprol}$ + $TU_{pmax}$ and $TU_{pprol}$ + $M2_{pkill}$ (right).

Dots represent average and error bars represent standard deviation of six technical replicates. Lines represent PCABM output with the median of optimized parameters. Shading represents model output for optimized parameters within interquartile range given by 50 optimizations.

### Supplementary Figure 6

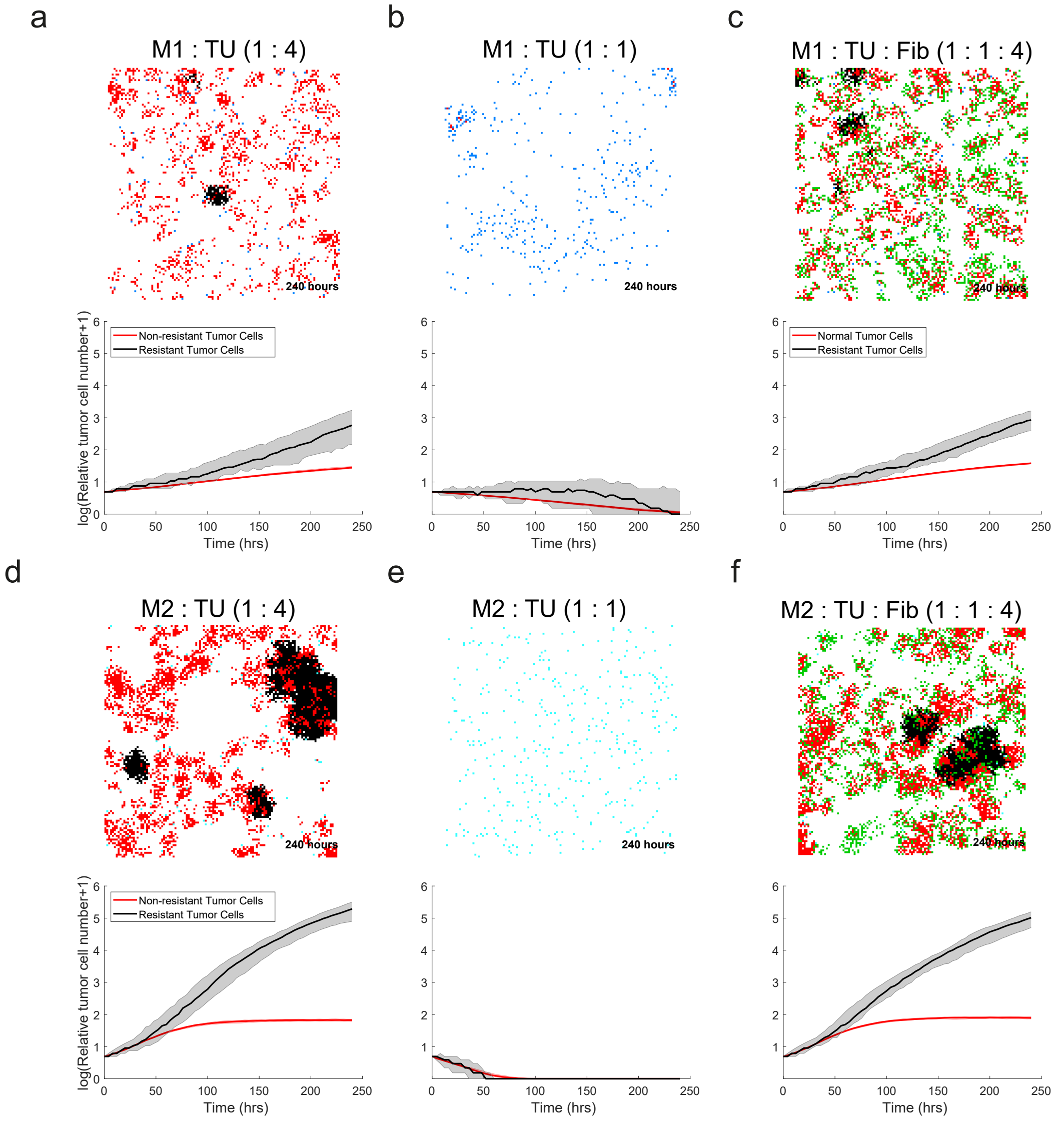

**Supplementary Figure 6: CRPC simulations**

**a)** LNCaP cells co-cultured with M1 macrophages at a 1:4 ratio

**b)** LNCaP cells co-cultured with M1 macrophages at a 1:1 ratio

**c)** LNCaP cells co-cultured with M1 macrophages and fibroblasts at a 1:4:4 ratio

**d)** LNCaP cells co-cultured with M2 macrophages at a 1:4 ratio

**e)** LNCaP cells co-cultured with M2 macrophages at a 1:1 ratio

**f)** LNCaP cells co-cultured with M2 macrophages and fibroblasts at a 1:4:4 ratio

LNCaP-abl cells are present in all simulations and are seeded with LNCaP cells at a 1:100 ratio. The initial amount of tumor cells does not change throughout the different simulations. Upper panels: LNCaP cells are shown in red, LNCaP-abl cells are shown in black, fibroblasts are shown in green and macrophages are shown in blue. Simulated growth after 240 hours (60 iterations) is shown.

Lower panels: lines and shading represent the median and interquartile range of 50 simulations. Red lines are relative growth for LNCaP cells and black lines are relative growth for LNCaP-abl cells.
